# Surgical techniques and tips for a reliable murine model of primary and metastatic pancreatic cancer

**DOI:** 10.1101/2022.07.30.502136

**Authors:** Jonathan J. Hue, Mehrdad Zarei, Hallie J. Graor, Omid Hajihassani, Erryk S. Katayama, Alexander W. Loftus, Ali Vaziri-Gohar, Jordan M. Winter

## Abstract

For patients with pancreatic cancer, survival rates lag behind other common cancers. This is in part due to the relative resistance to conventional chemotherapeutics and novel immune- or targeted-therapies. Ongoing research efforts are needed to identify and validate effective therapies. It is the unfortunate reality that a significant proportion of pre-clinical success does not translate into improved patient outcomes, likely due to a multitude of factors. In the current research landscape, flank xenograft models are commonly utilized to study pancreatic cancer, as this technique is fast, fairly non-invasive, and reliable. However, this model is not anatomically or physiologically accurate, does not impact other intra-abdominal organs, and experiments are often ended based on tumor size rather than systemic illness. Orthotopic injections of cancer cells directly into the pancreas for study of localized disease or into the spleen for study of hepatic metastases can be performed via a quick, reliable, minimally invasive surgical procedure with minimal morbidity and mortality. Existing methodologic reports are often sparse. Thus, there are significant knowledge and technical gaps for researchers attempting these techniques for the first time. In the current report, details of orthotopic pancreatic injections and splenic injections for metastatic disease are provided. Details of commonly encountered operative issues and mistakes are presented with suggestions to improve performance are described. A summary of expected outcomes is also provided herein.

## INTRODUCTION

Until more sensitive early detection methods are developed, it is the unfortunate reality that many patients with pancreatic cancer (PC) will present with advanced disease, not amenable to surgical resection (1–3). In this patient population, systemic therapies represent the only chance to lengthen survival. Patients with metastatic disease who forgo therapy typically survive on the order of months (4,5). Unfortunately, even with multiagent chemotherapy, most patients with metastatic disease often succumb within one year (6–8). A recent analysis by our research group demonstrated that the field of pancreatology is still at least several years away from development and approval of a novel, paradigm-shifting systemic therapy potentially leading to improved patient outcomes (9). Additionally, it is estimated that just 3% of drugs that enter clinical testing achieve approval by the Food and Drug Administration (9,10). Thus, ongoing pre-clinical research in this field is desperately needed to identify new and effective therapies.

For many murine models of cancer, flank xenografts are utilized. This method of tumor implantation is fast, non-invasive, reliable, and subcutaneous tumor volumes can be measured at frequent intervals (11). While these models would be anatomically appropriate for the study of cutaneous malignancies, such as melanoma, cancers of the pancreas and other intra-abdominal organs, may not be accurately represented with a subcutaneous model. Intra-abdominal tumors often have physiologic ramifications, such as organ-specific dysfunction (i.e., pancreatic insufficiency, hepatic failure, etc.), mass effect (i.e., pain, gastric outlet obstruction, bowel obstruction), local invasion or spread (i.e., carcinomatosis, development of ascites), and systemic signs of illness. These effects are not often observed in subcutaneous models. Several studies have also determined that the location of PC tumors (i.e., subcutaneous or pancreatic) leads to differences in growth, vascularity, collagen deposition, immune infiltrate, and response to immunotherapy. Alternatively, patient-derived tumor samples can be subcutaneously implanted or implanted into the pancreatic parenchyma for study of human malignancies (11). Utilization of patient-derived models requires implantation into immunocompromised mice, thus the role of the immune system and study of immunotherapies is limited. This model is also quite expensive to establish and labor intensive, with an extended duration from initial implantation to experimentation (11).

Some may argue that all of the aforementioned methods are “artificial” models of PC, as the early tumorigenesis does not mimic de novo tumor formation. Studies have utilized autochthonous or inducible models of murine PC (11–13). Unfortunately, these models can be quite expensive, require meticulous breeding and genotyping, or produce litters of varying sizes which makes experimental planning difficult. Additionally, these models develop countless tumors throughout the pancreas, which does not mimic human disease. These models are often exceptionally harsh, leading to rapid death, thereby making comparative survival studies difficult. Surveying the literature, autochthonous and inducible models are infrequently utilized relative to other murine models.

Orthotopic injections may represent a middle ground by recapitulating the appropriate tumor macro- and microenvironment, while overcoming the unpredictability and labor intensity of autochthonous models. They can be performed in immunocompetent mice to study murine PC (allograft), including the impact of the immune system, or immunocompromised mice to study human PC cell lines (xenograft). Additionally, large numbers of injections can be performed in a single day without a decrement in the tumor uptake rate, enabling multi-arm survival analyses to be appropriately powered. Current methodologic reports detailing orthotopic injections are often quite limited in description of operative techniques and presume the researcher has knowledge of surgical techniques (14–18). This leaves significant knowledge gaps for researchers who are new to this procedure. Protocols with an emphasis on proper surgical technique and how to remedy common intraoperative errors are needed. Herein, we provide a protocol for inoculation of pancreatic tumors via orthotopic pancreatic injections and hepatic metastases via splenic injections, to aid in the study of PC.

## MATERIALS AND METHODS

Mice were maintained in pathogen-free conditions within a dedicated animal facility. Mice were fed normal chow and had access to water ad libitum. In general mice should be at least 6 weeks of age to ensure the pancreas is an appropriate volume to minimize the risk of tumor cell leakage. Of note, the pancreas of different mouse breeds is variable in size and texture.

### Step 1 - Preparation of cells

1. Preparation of cells for orthotopic pancreatic injections
  1. Cells should be at approximately 70% confluent at the time of preparation to maximize tumor uptake.
  2. After cell detachment, count viable cells and transfer the total number of desired cells to a 15 mL conical. In general, prepare enough cells for approximately double the number of planned mice (i.e., prepare enough cells for 20 injections if 10 mice are required). Centrifuge at 4°C for 5 minutes at 1500 rpm. Aspirate supernatant and resuspend cell pellet in 5 mL of cold PBS.
  3. Centrifuge at 4°C for 5 minutes at 1500 rpm. Aspirate supernatant. Resuspend cell pellet in Matrigel followed by PBS (50-60% PBS, 40-50% Matrigel). The final volume should reflect the desired cell concentration for injection.
    1. Typically ~30 *μ*L is a safe volume to inject into the pancreas. Greater volumes increase the risk of leakage. Smaller volumes are difficult to accurately inject.
  4. Adequately mix the cell suspension. Transfer 1 mL of the cell suspension to a 1.5 mL Eppendorf tube to ensure adequate mixing is possible prior to injection. Keep cell suspension on ice until injection.
2. Preparation of cells for hepatic metastases via splenic injection
  1. Follow steps 1-2 as above.
  2. Resuspend cell pellet in cold PBS <u>without</u> Matrigel. The final volume should reflect the desired cell concentration for injection as above.
    1. Typically 100-200 *μ*L is injected into the spleen.
  3. Adequately mix the cell suspension. Transfer 1 mL of the cell suspension to a 1.5 mL Eppendorf tube. Keep cell suspension on ice until injection.

### Step 2 - Preparation of equipment and medications

1. Surgical instruments
  1. For orthotopic pancreatic injections, the following surgical instruments are recommended:
    1. One pair of fine tipped scissors
    2. Two pairs of blunt tipped, atraumatic forceps
    3. One needle driver
    4. One 4-0 permanent suture (e.g., Prolene or nylon)
    5. Three 27-gauge needles
    6. Three 1 mL syringes
    7. One wound closure device with skin clips
  2. For splenic injections, the following additional instruments are needed:
    1. One 4-0 silk suture (without attached needle)
    2. Sterile cotton swabs
  3. All instruments should be autoclaved prior to the procedure or left in sterilized single use packs (needles, syringes, suture). Note, if greater than 5 mice are undergoing injections at a time, multiple surgical packs are recommended.
2. Medications
  1. Isoflurane
  2. Oxygen
  3. Bupivacaine, lidocaine, or equivalent local analgesic
  4. Carprofen, meloxicam, or equivalent systemic analgesic
  5. Eye lubricant
3. Other equipment
  1. Betadine solution
  2. Alcohol swabs
  3. Electric hair shaver (if furred mice are being used)
  4. Two small rodent heating pads or reusable gel hot packs
  5. Glass bead sterilizer
  6. Anesthetic circuit including chamber and appropriately sized nose cone
  7. Clean cage for postoperative care
  8. Sterile gauze or drape

### Step 3 - Preoperative preparation

1. Operations should be performed on a clean surface, in a room with minimal foot traffic.
2. Turn on rodent heating pad and cover in water resistant disposable pad. Place a clean cage with easy access to food and water on a separate heating pad for mice to recover postoperatively.
3. Draw up local and systemic analgesics into separate labeled 1 mL syringes and attach 27-gauge needles.
4. Place mouse into anesthetic induction chamber until asleep. Transfer mouse to heating pad, place nose in nose cone, and position with left side up. Apply small amount of eye lubricant to both eyes.
5. Ensure adequate depth of anesthesia by performing a toe pinch. Adjust anesthetic flow rate if the mouse reacts.
6. Shave the left flank of mice (if furred) so that area just below the left ribcage is free of fur (approximately a 2 × 2 cm area), dispose of excess fur. If time permits, mice can be shaved prior to the procedure.
7. Apply betadine solution in a circular motion, starting from the desired incision site (just below the left rib cage; often the spleen can be visualized and can be used as a landmark) and working outward. Repeat two more times (total of three). Wipe off betadine with a sterile alcohol swab.
8. Surround operative site with sterile gauze or surgical drape
9. Open instrument packs and individual suture packs. Wash your hands and don sterile gloves.

### Step 4 - Orthotopic pancreatic injection

1. Inject local analgesia in the skin near the incision. Administer the systemic analgesic as directed (subcutaneous or intraperitoneal).
2. Using your non-dominant hand, gently grab the skin overlying the spleen (just underneath the ribcage) with blunt forceps and lift up, tenting the skin. Using your dominant hand, make a small incision using the tips of the scissors underneath where your forceps are holding, approximately 0.5 cm, in a transverse/horizontal direction
  1. The location of this incision is of paramount importance. An incision too superior (towards the mouse’s head) may result in inadvertent entry into the thoracic cavity, which is not well tolerated. An incision too inferior (towards the mouse’s feet) can lead to subsequent wound complication (i.e., the staple can easily be chewed out). An incision that is too anterior (towards the mouse’s stomach) can result in inadvertent injury to the liver, resulting in significant hemorrhage.
3. Gently grab the superior free edge of skin (closest to the mouse head) and lift upward. Gently insert the tips of your scissors into the incision and lightly spread several times to begin dissection through the subcutaneous tissues. Reposition your forceps and grab the exposed subcutaneous tissues (rather than the skin edge) and retract upwards. Continue working through the subcutaneous tissues, alternating between light spreading with the tips of scissors and cutting small pieces of tissue.
  1. Your scissors should be perpendicular to the body of the mouse (i.e., you are working your way into the abdominal cavity). Try to avoid tunneling upwards, as there is a risk of entering the thoracic cavity, which is not well tolerated.
4. Eventually, you will reach the abdominal wall musculature and peritoneal lining. This will feel “thicker” between your forceps. Make a small incision using your scissors and you should see into the abdominal cavity. Spread gently with your scissors to increase the length of the abdominal wall incision. This should match the length of your skin incision (~0.5 cm)
5. Gently grab the “top lip” of the abdominal wall with your non-dominant hand and retract upward. With your dominant hand, lightly grab the tail of the pancreas or spleen using blunt forceps and gently pull it out of the incision.
  1. In the authors’ experience, gently grabbing the tail of the pancreas is faster, more reliable, and has a reduced risk of a major complication (splenic hemorrhage).
6. Prepare your cellular mixture for injection. Gently invert the Eppendorf tube several tubes to ensure adequate mixing. Using the 1 mL syringe and 27-gauge needle, draw up 30 *μ*L of the cell suspension. Ensure the mixture is free from air bubbles prior to injection.
7. Lightly grab the tail of the pancreas with your non-dominant hand using blunt forceps and gently pull outward to “straighten the pancreas out.” Identify an appropriate injection site. Look for an area without obvious blood vessels and with enough surface area to minimize the risk of leaking. Be aware of “folds” in the pancreas, as injection here may result in multiple puncture sites.
8. Carefully insert the needle into the injection site with the bevel of the needle up, parallel to the long-axis of the pancreas. Traveling 3-5 mm through the parenchyma will minimize leakage, if possible. Slowly inject the cell mixture. You should see a small “bubble” form (**Figure 1**). After successful injection, slowly withdrawal the needle. Do not manipulate the pancreas for approximately 30 seconds to allow the injection site to seal to minimize leakage. Ensure there is no evidence of bleeding.
9. Using both pairs of blunt forceps, gently grab the top and bottom lips of the abdominal wall. Gentle “open and close” or rotational motions often allow the spleen and pancreatic tail to return to the abdominal cavity. Alternatively, you can gently guide the spleen back into the abdominal cavity. Do not grab the pancreatic tail near the injection site with forceps as this will cause tumor cell leakage.
10. Ensure the spleen and pancreas are completely returned to the abdominal cavity. Using the 4-0 permanent suture, suture the abdominal wall closed using a “figure-of-eight” method.
  1. Starting on the “bottom lip” on the right-side of the incision (towards the mouse back), drive the needle through the abdominal wall using the needle driver. The needle should travel from outside the abdominal cavity to inside the abdominal cavity.
    1. This suture should start approximately 1 mm from the right apex of the incision to ensure adequate closure. Ensure your needle tip does not puncture any underlying structures.
  2. Drive the needle through the “top lip” of the right side of the incision (1 mm from the right apex), this time from inside the abdominal cavity to outside the abdominal cavity.
  3. Drive the needle through the “bottom lip” of the left side of the incision (1 mm from the left apex) from outside the abdominal cavity to inside the abdominal cavity.
  4. Drive the needle through the “top lip” of the left side of the incision (1 mm from the left apex), from inside the abdominal cavity to outside the abdominal cavity.
  5. Gently pull up on the suture to close the wound and ensure that no intra-abdominal contents are in the incision. Pull the suture through so that there is approximately a 2 cm “short tail”. Perform an instrument tie to close the incision.
    1. With a needle driver in your dominant hand and the “long tail” of the suture (the side of the suture with the needle attached) in your non-dominant hand, wrap the suture around the needle driver. Open the jaws of the needle driver and grab the short tail of the suture. Pull the short tail of the suture through the loop of suture you created. With light upward tension, pull with your non-dominant hand until the incision is closed tight (to ensure intra-abdominal organs are not caught in your suture).
    2. Repeat step 3.10 three more times (total of four knots).
  6. Ensure the entire length of the abdominal wall incision is closed. Cut the suture tails directly on the knot of suture.
11. Close the skin incision with a wound clip. Grab the left-most corner of the skin incision and gently lift up. This should bring the skin edges together with good opposition. Often times, a single wound clip can be used to adequately close the incision, but multiple clips can be used as needed.
  1. Upward tension also serves to separate the skin from the underlying abdominal wall. Ensure only the skin is clipped, and not the abdominal wall as this can cause increased pain and wound complications.
12. Remove the mouse from the anesthetic circuit and place into a clean, dry cage. Monitor during its recovery until fully awake and moving.
13. If multiple mice are undergoing injections, place instruments into a glass bead sterilizer. Multiple “surgical packs” of instruments are recommended for large numbers of injections.

**Figure 1.**
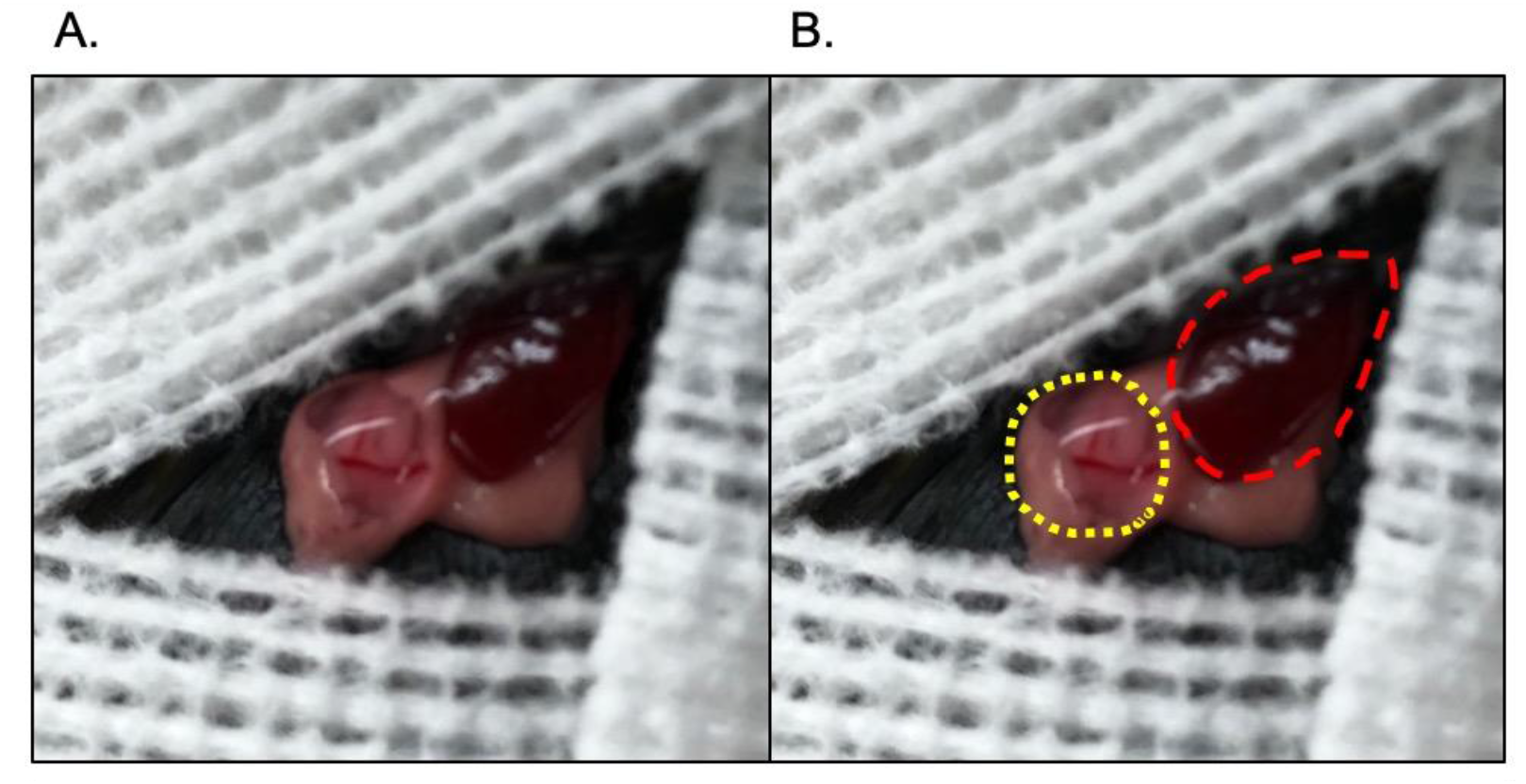
Intraoperative depiction of pancreatic tumor injection. A.) Still photograph. B.) Yellow dashes encircle the pancreatic injection “bubble,” red dashes encircle the spleen.

### Step 4 - Splenic injections

1. Follow steps 3.1 - 3.6 above to gain access to the abdominal cavity and externalize the spleen.
2. Carefully hold the spleen with blunt forceps in your non-dominant hand. With your dominant hand, carefully insert the needle ≥5 mm into the splenic parenchyma. Slowly inject the cellular mixture into the spleen, just underneath the capsule, while slowly withdrawing the needle. This will cause a “white-out” of the spleen.
3. After injection, quickly apply direct pressure to the injection site with a sterile cotton swab. Bleeding typically stops after 1-2 minutes, but may take longer depending on the depth of injection and trauma to the spleen. Wait for approximately 8-10 minutes. This is adequate time for cells to enter the splenic vasculature and travel to the liver via the splenic vein.
4. Perform a splenectomy by looping the 4-0 silk suture around the splenic hilum. Typically, a small amount of pancreas is incorporated into the suture. Tie three knots as was described above, ensuring the knots are tight and secure to prevent bleeding. Cut the tails of suture, leaving a 1-2 mm tail.
5. Using scissors, carefully cut through the splenic hilum taking care not to cut your suture, removing the spleen. Ensure that the splenic vessels are not bleeding. If bleeding, attempt to place another suture around the splenic vessels and tie tight.
6. Close the abdominal wall and skin, and recover the mouse as described above. Mice undergoing splenectomy often take longer to recover from anesthesia as compared to pancreatic injections due to the increased anesthetic time.

### Step 5 - Avoiding and fixing common mistakes

1. The cell injection leaked from the pancreatic injection site.
  1. This is by far the hardest part of the procedure. After grabbing a small piece of the pancreas, try to straighten it out. When injecting, match the angle and direction of your needle with the long axis of the pancreas to minimize the likelihood of going “through and through.”
  2. Each mouse pancreas is different: try to be flexible in your injection method and location. Sometimes the needle should be advanced several millimeters into a more suitable area, while other times the best location is directly under your supporting forceps. Don’t attempt an injection unless you are confident. Each unnecessary needle hole increases the likelihood of leakage.
  3. Very slowly begin to inject and closely monitor the injection site. If leakage is noted, stop the injection and attempt a separate injection site at a different location, far from the first site. Attempt to absorb the leaked cell suspension using sterile gauze.
  4. Practice prior to the planned experiment. After 15-20 orthotopic injections, the proportion of injections which leak significantly will decrease drastically in the author’s experience.
2. The spleen started to bleed while attempting to externalize it.
  1. The spleen will bleed significantly if the capsule is disrupted, so great care must be used. In general, it is safer to grab a “bigger piece” of the spleen, rather than a “smaller piece” as the force of the forceps is distributed over a larger area.
  2. Attempt to hold light pressure with a sterile cotton swab. Try to keep the spleen externalized, otherwise holding pressure is difficult. If bleeding cannot be controlled, a splenectomy can be performed (as described in Step 4).
  3. Lightly grabbing the pancreas, instead of the spleen, is safer in the authors’ opinion.

4. The abdominal wall is difficult to close.
  1. This can occur for a variety of reasons, including poorly placed initial skin incisions requiring the abdominal wall incision to be extended or “jagged” appearing incisions due to multiple cuts. Often times, driving the needle through healthy appearing tissue near the incision apices will adequately close even large or jagged incisions. Attempt to have the abdominal wall evert, rather than invert, to facilitate healing.
  2. If a single figure-of-eight suture does not completely close the incision, a second suture can be placed.
5. The skin incision is difficult to close.
  1. This is fairly common, often due to asymmetric dissection through the subcutaneous tissues. Prior to deploying the skin clip, the skin edges should be able to move freely without the underlying musculature to facilitate appropriate healing.
  2. Blunt tipped forceps can be used to free the skin edges from the underlying attachments of the musculature. Alternatively, scissors can be used to cut small tissue “bridges.”
  3. The skin edges should be everted, rather than inverted to facilitate healing
6. The skin clip fell out or was removed by the mouse prior to complete wound closure.
  1. This is not uncommon, but the risk can be minimized by a well-placed incision. If the clip is removed early and abdominal wall suture remains intact, the incision can be washed out and re-closed. If the clip is removed later in the postoperative course, often these mice will recover without issues. Additionally, antibiotic ointment can be applied to facilitate healing.
  2. On rare circumstances, both the skin clip and underlying suture fail. It is not recommended to attempt re-closure, as the peritoneal cavity is likely contaminated. Unfortunately, these mice often require euthanasia.

## RESULTS

Author JJH has performed greater than 1200 orthotopic pancreatic injections to date. The operative mortality rate is 0%, meaning that all mice tolerated surgery well and awoke from anesthesia without issue. Using murine PC cells in immunocompetent C57BL/6J mice, tumors form in approximately 80-90% of injected mice, even if the cell suspension injection is complicated by a small amount of leakage (**Figure 1A-B**). The tumor formation rate is similar in immunocompromised athymic nude mice injected with human PC cell lines. The injection of murine PC cells into immunocompromised mice (athymic nude or NOD scid gamma (NSG)) has yielded tumors in 100% of mice to date (N=80). The cells utilized by our research team express luciferase, thus viable tumors can be confirmed using bioluminescence imaging after intraperitoneal injection of luciferin (**Figure 2A-B**). Bioluminescence imaging can also be used to monitor growth over time (**Figure 2C**), similar to external measurement of subcutaneous xenografts. Additionally, this imaging modality can be used to confirm hepatic metastases after splenic injection (**Figure 2D**).

**Figure 2.**
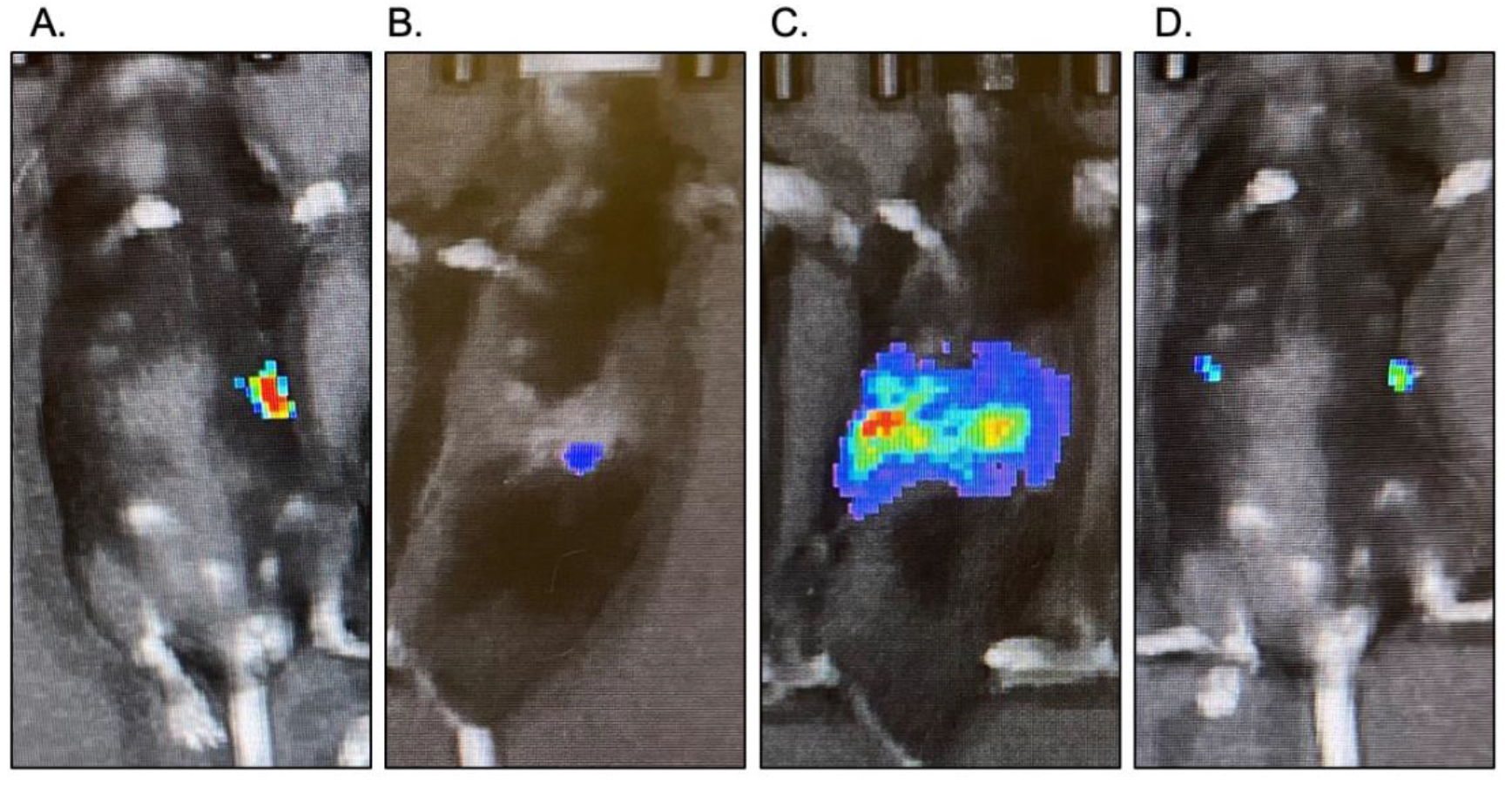
Tumor visualization using bioluminescence imaging. A-B.) Tumor validation 7 days after orthotopic pancreatic tumor implantation (supine and left side up). C.) Evidence of significant tumor growth 45 days after tumor implantation. D.) Validation of simultaneous orthotopic pancreatic tumor implantation (left-side of mouse) and hepatic metastatic tumor (right-side of mouse).

Based on our experience, median survival after orthotopic pancreatic injections is related to the number of cells injected. For example, median survival after injection of 10,000 murine PC cells is approximately 60-70 days from tumor implantation. Injection of 30,000-50,000 cells leads to a median survival of approximately 40-50 days. Thus, the cell number can be manipulated based on the researcher’s experimental timeline. Mice with confirmed orthotopic pancreatic tumors develop large tumors over the course of several weeks (**Figure 3A**). Mice also develop malignant ascites with large pancreatic tail tumors (**Figure 3B-C**). Mice only develop ascites late in the disease course and do not have evidence of widespread carcinomatosis, suggesting this is unlikely to be from cell leakage at the time of initial injection.

**Figure 3.**
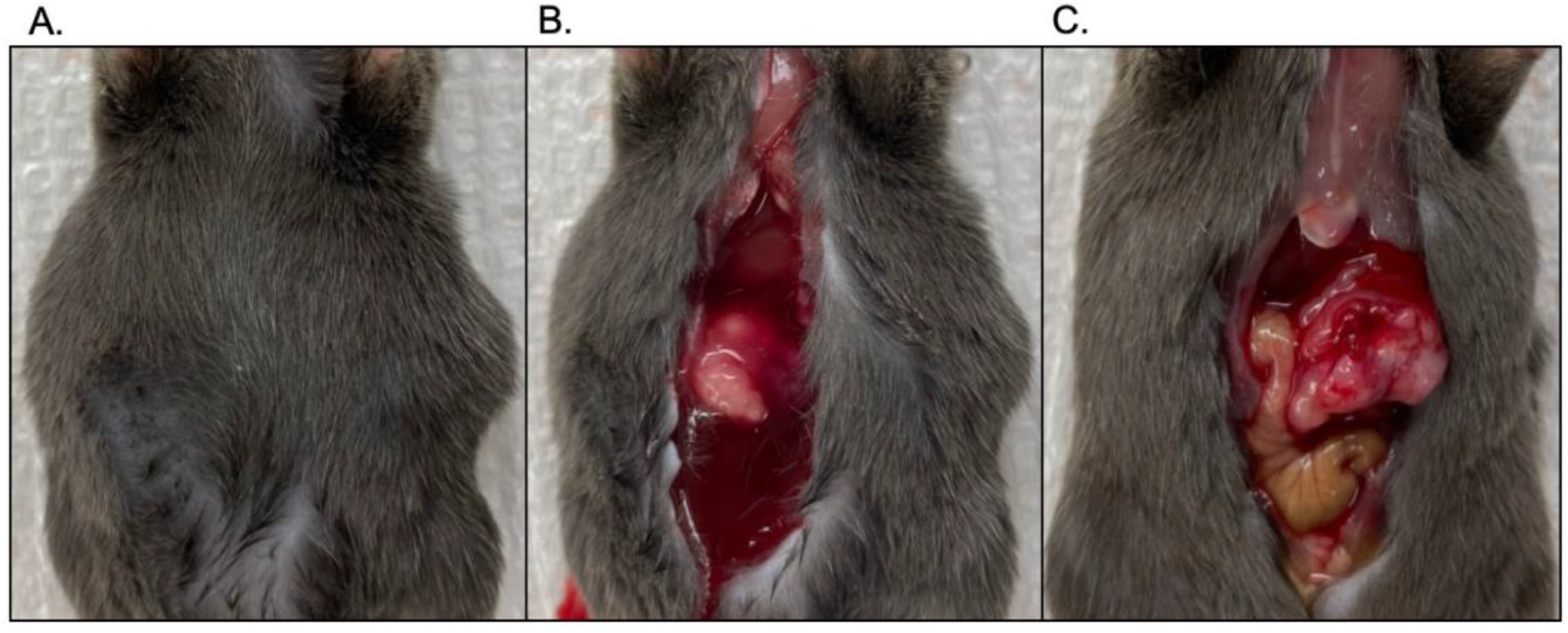
Evidence of intra-abdominal sequela in a mouse with an orthotopic pancreatic tumor. A.) Significant abdominal distension is noted upon external inspection. B.) Upon exploration, significant bloody ascites is identified. C.) A large tumor in the tail of the pancreas is identified.

Immunocompetent mice that underwent orthotopic pancreatic injection with simultaneous splenic injection develop a single primary pancreatic tumor and multi-focal hepatic metastases (**Figure 4A-B**). Interestingly, immunocompromised NSG mice develop spontaneous hepatic metastases after orthotopic pancreatic injections (i.e., without splenic injection of tumor cells; **Figure 4C-D**). The gross examination of these specimens suggests the splenic injection of tumor cells closely recapitulates the development of spontaneous metastases.

**Figure 4.**
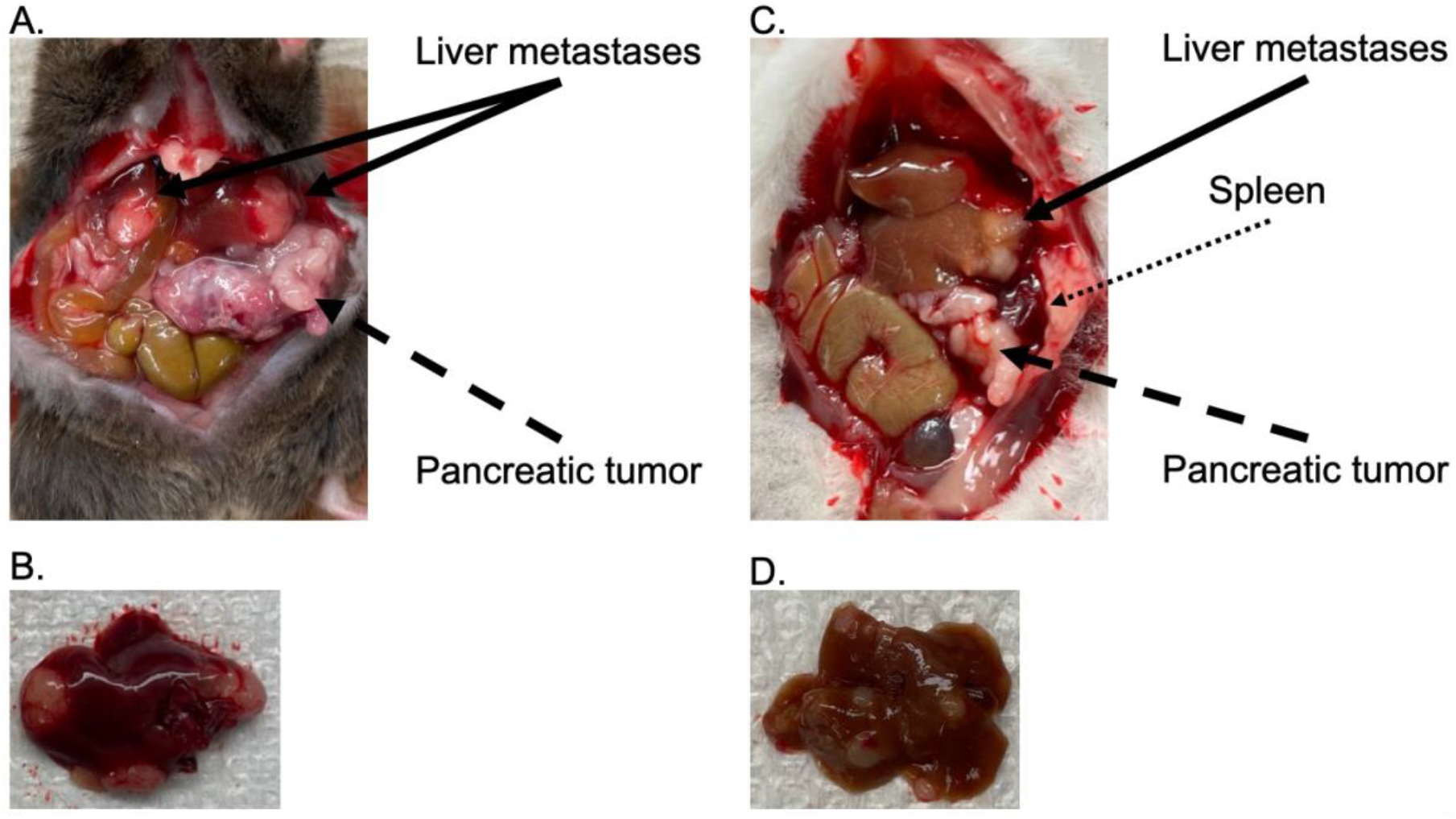
Orthotopic splenic injections in immunocompetent mice yield liver tumors similar in appearance to spontaneous liver metastases in immunocompromised mice. A-B.) Evidence of pancreatic tumor with hepatic metastases in C57BL/6J mice following simultaneous orthotopic pancreatic and splenic injections. C-D.) Evidence of pancreatic tumor with spontaneous hepatic metastases in NSG mice following orthotopic pancreatic injection (without splenic injection).

Lastly, tumor growth and appearance vary dependent on the location of the implanted PC cells (**Figure 5**). A lesser number of cancer cells injected into the pancreas grow larger than cells injected subcutaneously after one month. This finding is similar to past reports (19).

**Figure 5.**
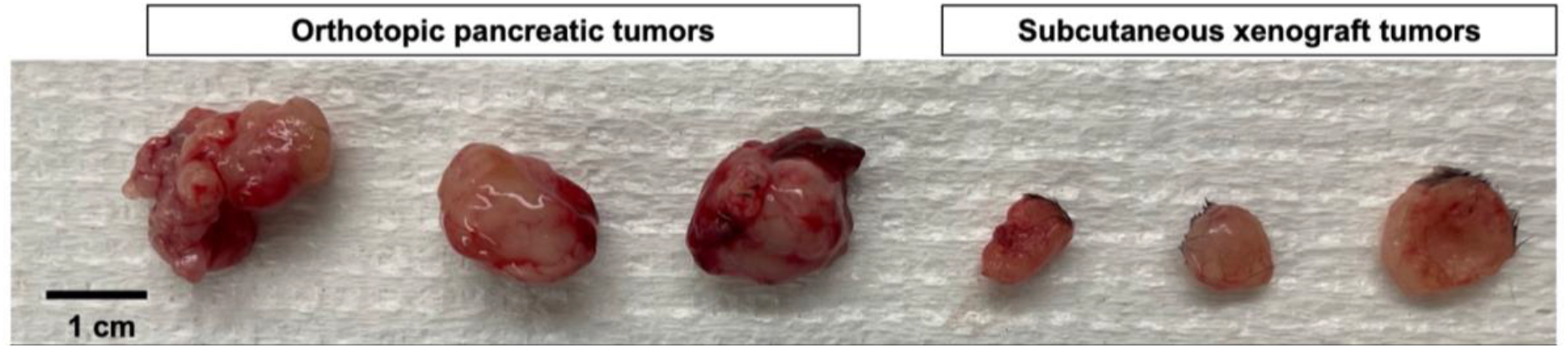
Growth of murine pancreatic cancer cells is variable based on tumor location. Murine PC cells were injected into the tail of the pancreas (15,000 cells) or subcutaneous tissue (60,000 cells). All tumors were harvested after 30 days of growth.

## DISCUSSION

Herein we present step-by-step instructions for development of primary and metastatic pancreatic tumors. The existing literature on these methods leaves gaps in technical steps and knowledge. Widespread adoption of these procedures may narrow the gap between pre-clinical and clinical results. As an example, FOLFIRINOX (folinic acid, 5-fluorouracil, irinotecan, oxaliplatin) was modestly effective in treating orthotopic pancreatic tumors but virtually ineffective for flank tumors (19). FOLFIRINOX is currently the most effective systemic therapy approved for treatment of PC (20–22), thus the lack of efficacy in flank tumors demonstrates the importance of tumor location. Additionally, recent studies have shown that subcutaneous murine PC tumors respond well to certain checkpoint inhibitors (23–26). However, orthotopic tumors were shown to be resistant to checkpoint blockade (26–28), which may be more akin to patients with PC (29).

Currently, the majority of patients with PC present with metastatic disease, of which the liver is the most common site of extra-pancreatic disease. However, relatively few pre-clinical studies investigate the efficacy of novel therapeutics in mice bearing hepatic metastases. It is well documented that primary and metastatic tumors have unique tumor microenvironments, likely causing variable responses to therapy. More widespread adoption of simultaneous pancreatic and splenic injections may identify unique treatment combinations with efficacy in both tumor deposits.

The surgical procedures described have several key steps. Perhaps most importantly, is maintaining animal welfare by ensuring appropriate depth of anesthesia throughout the procedure, meticulous surgical technique, and adequate analgesia in the perioperative period. Proper tissue handling will limit postoperative pain and yield superior results. From a technical standpoint, delicate handling of the spleen to prevent capsule violation is crucial. Severe bleeding from a ruptured spleen is difficult to stop and may require an unplanned splenectomy. Injection of pancreatic cells into the pancreas takes great precision and practice. The optimal pancreatic injection site is different for each mouse and care must be taken to avoid unnecessary needle holes in the pancreatic capsule. This procedure can be modified to fit the needs of individual researchers. The easiest variable to manipulate is the number of cells injected, as a greater number of cells often results in earlier mortality. Lower cell numbers may theoretically recapitulate de novo PC development, but risks a lower rates of successful tumor uptake. Additionally, these procedures may also be performed using human PC cell lines in immunocompromised mice. Typically, a greater number of cells (100,000 - 500,000) are required to maximize tumor uptake.

While the surgical injections described in this protocol are likely an improvement from commonly used subcutaneous xenograft models, it is not without limitations. Notably, this is still an artificial model. The autochthonous model of PC development likely represents a model more akin to human disease progression. Development of an autochthonous model is a time consuming and arduous process, which requires extensive mouse breeding, frequent genotyping, and leads to relatively unpredictable sample sizes. Performing experiments with adequate sample sizes to detect statistical significance with the autochthonous model is difficult. Further, mice develop multi-focal pancreatic tumors leading to rapid death. Using the methods described herein, exact numbers of immunocompetent mice can be injected with high rates of successful tumor uptake.

Orthotopic implantation of pancreatic and hepatic metastatic tumors represent a model which is fairly similar to normal physiology and feasible to perform in large sample sizes. While flank xenograft tumors are fairly non-traumatic and technically easy to perform, they lack the ability to cause intra-abdominal sequela, often seen in patients with PC. Additionally, preliminary data from our research lab suggests that the cellular makeup of xenograft and orthotopic tumors are significantly different. As an alternative to splenic injections, researchers commonly perform tail vein injections for development of a fairly non-invasive metastatic model. The tail vein drains directly into the inferior vena cava, thus almost exclusively leads to development of pulmonary metastatic deposits. However, just 4% of patients with metastatic pancreatic cancer present with pulmonary metastases, while 40% present with hepatic deposits and 55% present with peritoneal disease (30). Although these fairly non-invasive methods (subcutaneous and tail vein injection) are easy on researchers and mice, they do not accurately represent PC. The surgical procedures described in this protocol can be performed with a small incision (~0.5 cm) and completed in just several minutes in experienced hands.

## CONCLUSION

As PC is often described as the deadliest of common cancers, novel therapies are desperately needed to improve patient outcomes. Widespread adoption of more physiologically appropriate models may better recapitulate the disease process. We hope this report will enable a greater proportion of PC researchers to include orthotopic models in their research endeavors.

## ACKNOWLEDGEMENTS

Thee authors would like to acknowledge Dr. Darren Carpizo for generously providing K8484 KPC cells.

## AUTHOR CONTRIBUTIONS

- Conceptualization: JJH, MZ, JMW
- Methodology: JJH
- Data Curation: JJH, MZ, HJG, AVZ
- Writing – Original Draft Preparation: JJH
- Writing – Review & Editing: MZ, HJG, OH, ESK, ALW, AVZ, JMW
- Supervision: JMW

## INSTITUTIONAL STATEMENT

All experiments involving mice were approved by the Case Western Reserve University Institutional Animal Care Regulations and Use Committee (IACUC, protocol 2018-0063).

